# Cortical inactivation does not block response enhancement in the superior colliculus

**DOI:** 10.1101/2020.03.19.998807

**Authors:** Katarzyna Kordecka, Andrzej T. Foik, Agnieszka Wierzbicka, Wioletta J. Waleszczyk

**Affiliations:** Nencki Institute of Experimental Biology, Polish Academy of Sciences, 3 Pasteur St., 02-093 Warsaw, Poland; International Centre for Translational Eye Research, Institute of Physical Chemistry, Polish Academy of Sciences, 44/52 Marcina Kasprzaka St., 01-224 Warsaw, Poland

**Keywords:** visual evoked potential, plasticity, repetitive visual training, visual cortex, superior colliculus, electrophysiology

## Abstract

Repetitive visual stimulation is successfully used in a study on the visual evoked potential (VEP) plasticity in the visual system in mammals. Practicing visual tasks or repeated exposure to sensory stimuli can induce neuronal network changes in the cortical circuits and improve the perception of these stimuli. However little is known about the effect of visual training at the subcortical level. In the present study, we extend the knowledge showing positive results of this training in the rat’s superior colliculus (SC). In electrophysiological experiments, we showed that a single training session lasting several hours induces a response enhancement both in the primary visual cortex (V1) and in the SC. Further, we tested if collicular responses will be enhanced without V1 input. For this reason, we inactivated the V1 by applying xylocaine solution onto the cortical surface during visual training. Our results revealed that SC’s response enhancement was present even without V1 inputs and showed no difference in amplitude comparing to VEPs enhancement while the V1 was active. These data suggest that the visual system plasticity and facilitation can develop independently but simultaneously in different parts of the visual system.

## Introduction

Repetitive visual training is a rapidly developing tool to modulate neuronal plasticity in the visual system for both research and clinical application [1]. Appropriate visual training protocols can strengthen residual vision in patients with visual impairments such as glaucoma [2], optic nerve neuropathy [3] or hemianopia [1,4,5].

Simultaneously repetitive visual training is successfully used in studies on the visually evoked potential (VEP) plasticity in the visual system of mammals. Repeated exposure to sensory stimuli can induce neuronal plasticity and leads to an enhanced visual response of these stimuli. Numerous researches have shown that the VEP enhancement might reflect synaptic plasticity [6–12]. A few days long repeated presentation of gratings with single orientation resulted in a potentiation of the cortical VEPs amplitude to the presented stimuli [9]. The mechanism underlying this form of training-dependent plasticity is known as long-term potentiation (LTP) of the cortical response [9,13,14]. There are well-described types of rapid VEP plasticity evoked by a few minutes of ‘photic tetanus’ stimulation [15]. The aforementioned repetitive stimulation results in positive changes in the visual system, for example, an expansion of neuronal receptive fields into unresponsive regions of the visual field [16]. The enhancement of the cortical response is an effect of sensory LTP dependent on NMDA-receptors in animals [15] and humans [10,17,18]. Studies on humans showed that repeated exposure to a checkerboard reversal stimulation leads to an increase of cortical VEP amplitude [8,19,20].

Little is known about the effect of visual training at the subcortical level. A study carried out on the rat’s dorsal lateral geniculate nucleus (dLGN), the primary recipient of visual information, suggests that the response properties of thalamic neurons are subject to experience-dependent long-term plasticity [21,22]. Similarly, in the superior colliculus (SC), the second major target of retinal input, repetitive exposure to dimming stimuli effectively induced the LTP of developing retinotectal synapses in *Xenopus* tadpole [23]. These reports suggest that appropriate sensory stimulation may induce plasticity also at the subcortical level. To test this hypothesis we used a 3 hours repetitive visual training to induce and determine the VEP plasticity in the primary visual cortex (V1) and the SC. Our study showed that visual training evokes enhancement of visual responses both, at the cortical and subcortical levels. Further, we revealed that repetitive visual training evokes response enhancement in the SC even if cortical input is turned off by xylocaine.

## Materials and Methods

### Subjects

We used 14 adults (200-250 g) male Wistar rats obtained from the Mossakowski Medical Research Centre. All animals were housed in the Animal House of the Nencki Institute of Experimental Biology and maintained on a 12 h light/dark cycle (light on 7:00 am). All experimental procedures were conducted in accordance with the *ARVO Statement for the Use of Animals in Ophthalmic and Vision Research* and the EC Directive 86/609/EEC for animal experiments using protocols and methods accepted by the First Warsaw Local Ethical Commission for Animal Experimentation (521/2018).

### Surgical procedures

Rats were deeply anesthetized with urethane (1.5 g/kg, Sigma-Aldrich, Germany, administered intraperitoneally) and placed in a stereotaxic apparatus. Additional doses of urethane (0.15 g/kg) were administered when necessary. Body temperature was maintained between 36 - 38°C using a heating blanket (Harvard Apparatus, MA, USA). Every hour fluid requirements were fulfilled by subcutaneous injection of 0.9 % NaCl. The skin on the head was disinfected with iodine and local anesthetic (1% lidocaine hydrochloride; Polfa Warszawa S.A, Poland) was injected. The craniotomy was done above the binocular V1 (6.5 - 7 mm posterior to Bregma; 4.5 mm lateral; [24]) and the SC contralateral to the stimulated eye (7.0 mm posterior to Bregma; 1.5 mm lateral; [24]. During recording the right eye (not stimulated) was covered with black tape. The Vidisic gel (Polfa Warszawa S.A, Poland) was applied to prevent the cornea from drying.

### Local field potential recordings and visual stimulation

Local field potentials (LFPs) were collected using linear electrodes made of 25 μm tungsten microwire in HML (Heavy Polyimide) insulation (California Fine Wire, US). The ground-reference electrode (Ag/AgCl wire) was located in the neck muscles. The cortical electrode (8 channels) was made with an inter-channel distance ranging from 100 - 300 μm and inserted 1.8 mm below the dura, passing through all cortical layers (supragranular, granular and infragranular). The SC electrode was located in the upper layers; stratum griseum superficiale – SGS, stratum opticum – SO. The SC electrode consisted of seven wires with a ∼ 200 μm vertical recording site arrangement and inserted 4 mm (tip) below the cortical surface. Signals were recorded with a multichannel data acquisition system (USB-ME64-System, Multichannel Systems, Germany), amplified 100 times (USB-ME-PGA, Multichannel Systems, Germany), filtered at 0.1–100 Hz, digitized (1 kHz sampling rate) and stored on the computer for offline analysis. Visual stimulation was controlled by Spike2 software (Cambridge Electronic Design, UK). Stimulation marks were recorded along with the electrophysiological signals. Flash VEPs were evoked using light-emitting diodes (LEDs, 2200 lx) positioned 15 cm in front of the rat’s left eye. The repeated visual training consisted of 300 flashes at 0.5 Hz repeated every 15 min for 3 hours. (Fig. 1A; [25]). Control recordings were carried out (100 flash repetitions at 0.1 Hz) before and after visual training to investigate the effect of training. A schematic diagram of the experimental protocol is shown in Figure 1.

**Figure 1.**
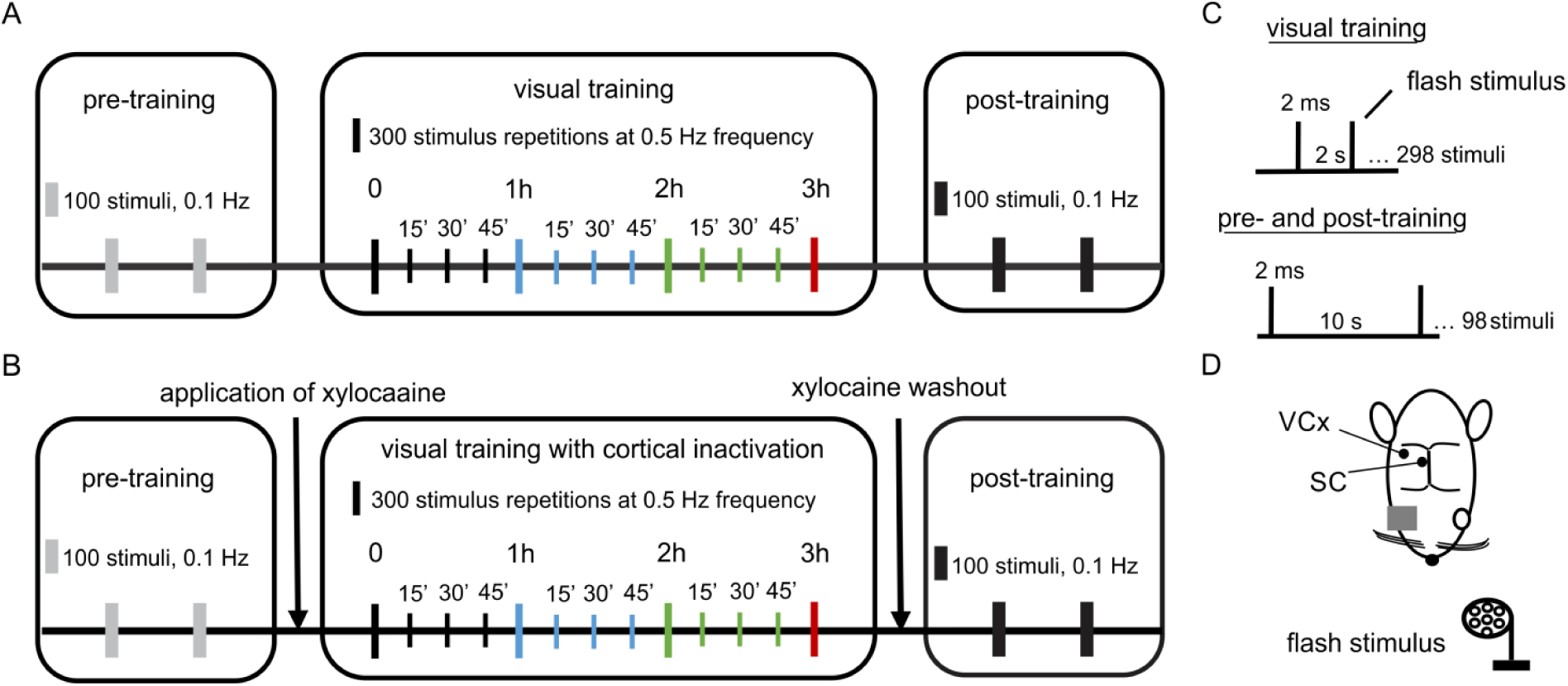
Design of an experiment revealing visual system plasticity caused by visual training. A, The visual training paradigm: 100 repetitions of light flashes at 0.1 Hz during pre- and post-training controls and 300 repetitions of light flashes every 15 min through 3 hours at 0.5 Hz during visual training. B, Visual training paradigm paired with cortical inactivation. The graph presents the same parameters of visual stimuli as in A. C, Visual stimulus parameters used in training. D, Schematic representation of the top view of the rat head with marked electrodes locations for the SC and the V1, both contralateral to the stimulated eye.

### Temporal inactivation of the cortex

A plastic chamber placed above the right hemisphere was filled with a xylocaine solution (2.5%, Lidocainum Hydrochloricum WZF, Polfa Warszawa S.A, Poland) to inactivate cortical activity (action potential blockage) during visual training [26]. The solution was replaced every 30 min during electrophysiological recordings.

### Data processing

Collected data (7 – visual training, 7 - visual training with cortical inactivation) was analyzed using Matlab (Mathworks). The LFP signal was preprocessed using the 1st order band-stop Butterworth filter at 50 Hz, a high-pass filter at 0.1 Hz, and a low-pass filter at 100 Hz. Continuous signals were divided into trials 1.2 s long (from 0.2 s before to 1 s after stimulus). The peak-to-peak amplitude of VEPs was calculated at every hour during visual training and control recordings in the 0 – 0.2 s time range, where 0 is a stimulus onset time. Cortical LFPs were analyzed in the delta (1-4 Hz) and theta (4-7 Hz) frequency ranges to monitor brain state. The mean frequency of the EEG signal occurring during recording was estimated at 1-5.6 Hz level. The signal to noise ratio was obtained by dividing the VEP amplitude by the peak to peak amplitude of the raw signal (%). For statistical analysis, two recording channels for each layer from a given structure were taken in each animal. The VEPs amplitudes and signal to noise ratio factor of pre- and post-training recordings were compared using two-tailed paired *t-*tests. To compare mean collicular VEP amplitude between the two experimental paradigms: V1 activated and inactivated, the two-tailed unpaired *t-*tests with Welch correction was used. One-way repeated-measures ANOVA with greenhouse Geisser correction was used with the Dunnett’s post hoc test to investigate changes in the mean VEPs responses and signal to noise ratio for 3 h of visual training. Results are presented as an average percentage of the mean response (± SEM) of the pre-training recording or the first session of visual training (time 0).

### Histology

Electrodes coated with DiI (1,I’-dioctadecyl-3.3,3’,3’ tetramethyl-indocarbocyanine perchlorate; Sigma-Aldrich, Germany) were used to facilitate electrode tract reconstruction [27]. At the end of the experiments, rats were given an overdose of Nembutal (150 mg/kg) and transcardially perfused with 4% paraformaldehyde in 0.1 M PBS (Sigma-Aldrich, Germany). The brains were removed, postfixed for 24 h in 4% paraformaldehyde, stored successively in 10%, 20% and then in 30% sucrose in 0.1 M PBS before sectioning. Brains were cut into 40 µm slices and stained with cresyl violet. The data from the experiments with incorrect electrode placements were excluded from further data analysis.

## Results

Analysis of VEP amplitude for the V1 included the granular layer (400-800 µm), with a typical reversal of potential polarity and infragranular layers (1000-1800 µm), identified by negative components. We considered layer V and VI as infragranular layers. We analyzed also the two retino-recipient layers of the SC: SGS (2.8 – 3.4 mm) and SO (3.6 – 4 mm), which were characterized by a biphasic oscillatory pattern of the recorded signal.

### Effect of visual training on the magnitude of VEPs amplitude

To test the effect of repetitive visual training we compared pre- and post-training responses to light flashes. Examples of cortical and collicular VEPs from a single rat are shown in Figure 2A-D. The comparison of pre- and post-training VEPs showed a larger response magnitude after 3 hours training session (see Methods for more details), both in the cortex and the SC (Fig. 2I-L). The stronger enhancement occurred in the SC compared to the V1. In the SGS layers of the SC, the amplitude of the post-training response was significantly greater (180 ± 9 %, n = 10, p < 0.001) than the pre-training amplitude. The increase in the SO layers was even stronger (208 ± 17 %, n = 10, p = 0.004). Results for both layers in the V1 revealed a significant increase of the VEP amplitude after visual training. The potentiated response in the granular layer after visual training was higher (179 ± 18 %, n = 12, p = 0.005) than the increase seen in the infragranular layers (144 ± 10 %, n = 14, p = 0.003).

**Figure 2.**
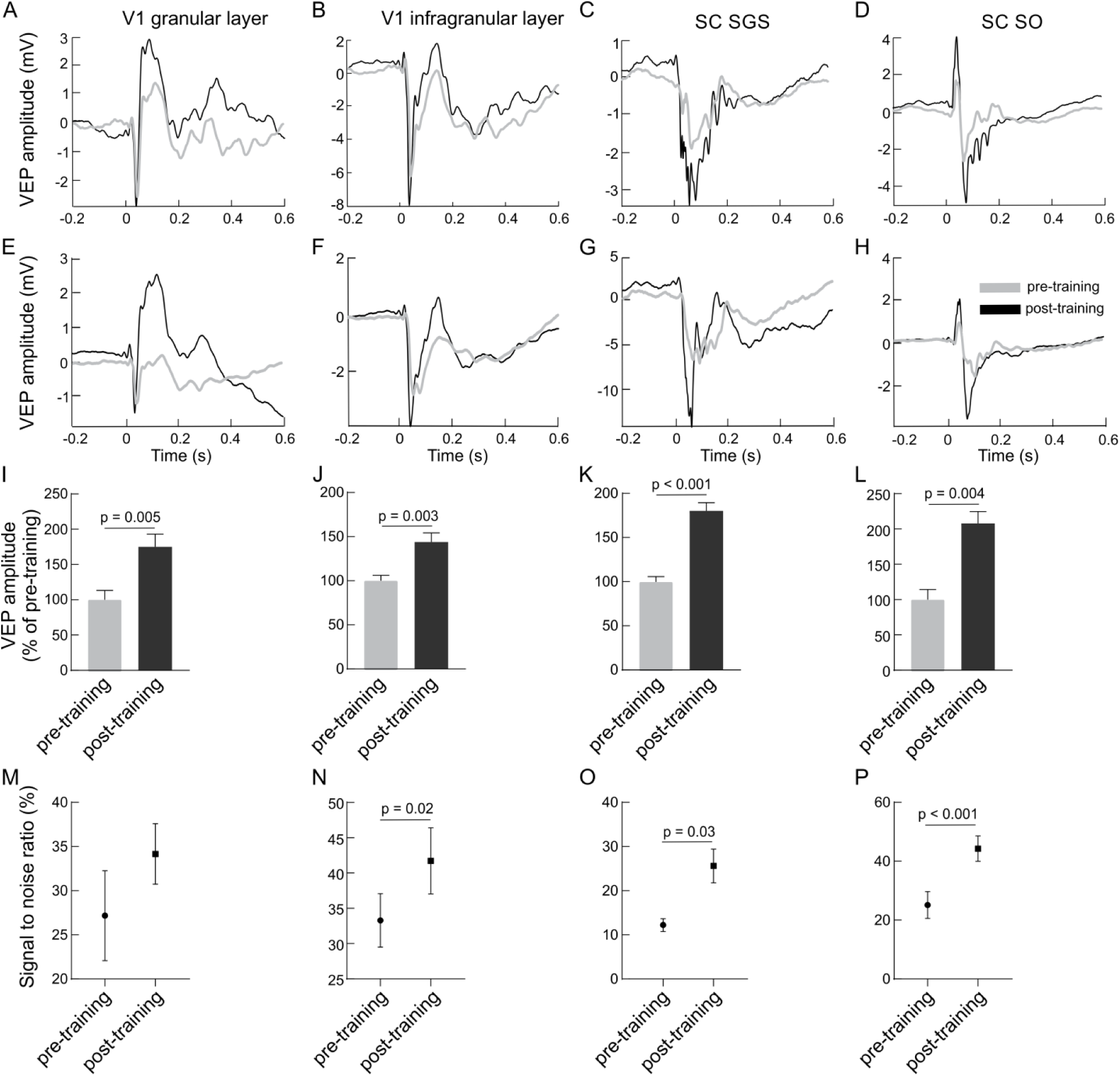
Visual training enhances visually evoked responses in the V1 and the SC. The first column presents results for a granular layer of the V1, second column for infragranular layers for the V1, the third column for the SGS layer and the last column for the SO layer of the SC. A-D, Average VEPs (n = 100) recorded in the V1 and SC for a single experiment before (grey line) and after (black line) visual training. E-H, Average responses in the V1 and SC for all rats before and after visual training. Time 0 is a stimulus onset. I-L, Comparison between the average pre-(grey bars) and post-training (black bars) VEP amplitudes for the V1 and SC. M-P, Signal to noise ratio changes for pre- and post-training recordings for the V1 and SC.

Figure 3 presents VEPs obtained for every hour of visual training in the V1 and the SC including division into layers. The largest increase of VEP amplitudes was observed after three hours of stimulation in all studied cases. Response in granular layer (Fig. 3I; F(3,11) = 4.76, p = 0.009) was significantly greater after 2^nd^ hour (183 ± 17 %, p = 0.04) and 3^rd^ hour of training (215 ± 33 %, p = 0.04) in comparison to control. In the infragranular layers (Fig. 3J; F(3,13) = 4.76, p = 0.0006) the greatest amplitude was during 3^rd^ hour (163 ± 8 %, p = 0.0005) as well as 2^nd^ hour of training (138 ± 6 %, p = 0.009) than in control.

**Figure 3.**
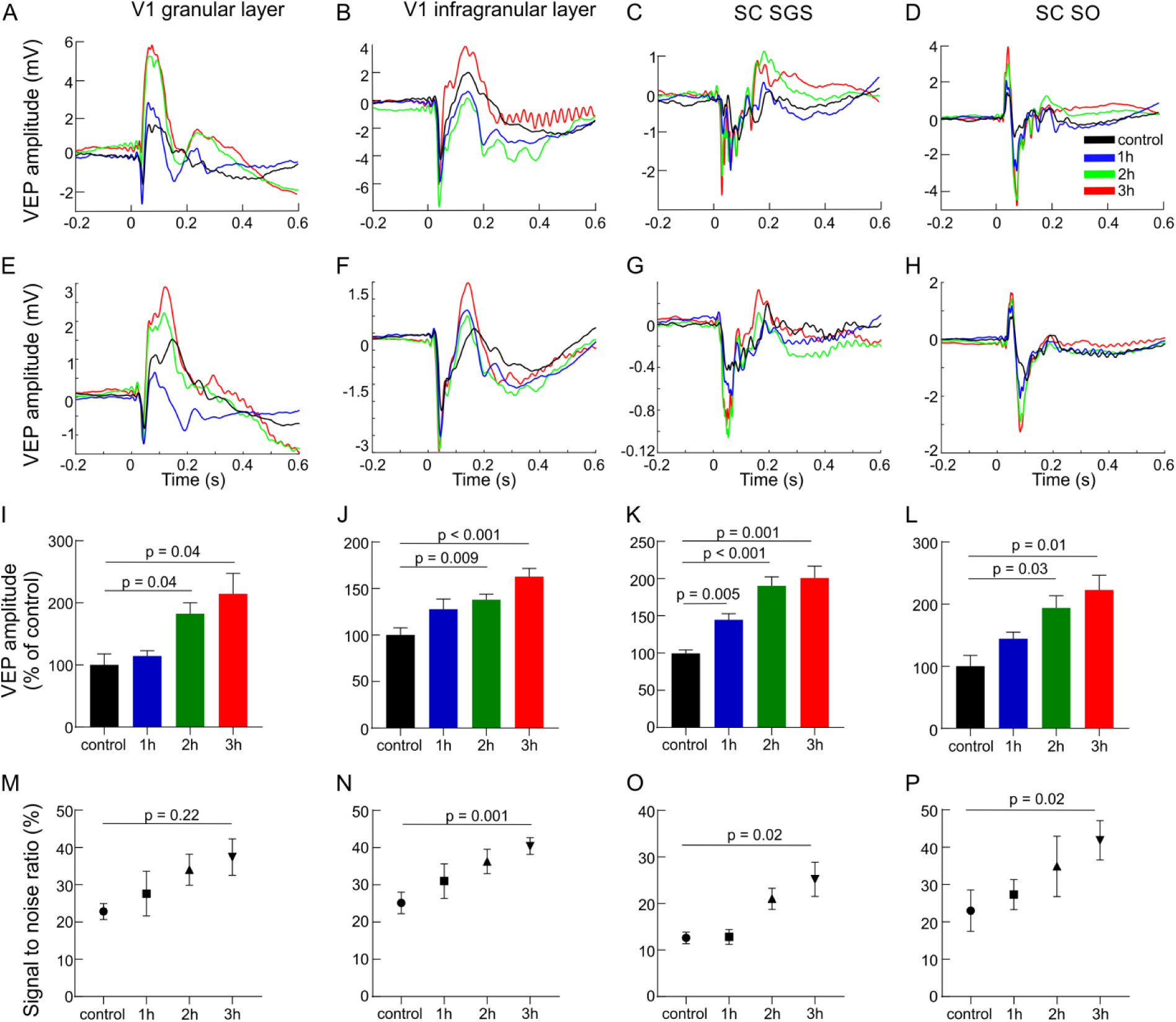
Response enhancement during visual training. The data are presented in order as in Figure 2. A-D, VEPs for each hour of visual training recorded in different layers of the V1 and the SC for a single experiment. E-H, Average population responses for each hour (every fourth sensory stimulation) during visual training recorded in different layers of the V1 and the SC and encoded by different colors. I-L, Mean VEP amplitudes for all rats and every hour of visual training for the V1 and SC. M-P, Signal to noise ratio changes for every hour of visual training for all layers.

Also in the SC, the visual training evoked significant VEP potentiation (for SGS; F(3,9) = 6.59, p < 0.0001, for SO; F(3,9) = 4.76, p = 0.005). In the SGS (Fig. 3K) the response was the highest after 3^rd^ hour of the training (201 ± 18 %, p = 0.001), then 2^nd^ hour (190 ± 14 %, p = 0.0005) and 1^st^ hour (145 ± 9 %, p = 0.005) in comparison to control. The same tendency was observed in the SO layer (Fig. 3L), the greatest visual response was after 3^rd^ hour of training (223 ± 24 %, p = 0.01), then 2^nd^ hour (194 ± 20 %, p = 0.03) and 1^st^ hour (144 ± 10 %, p = 0.2), however not significant.

### Effect of V1 inactivation on visual training efficiency in the SC

The potentiation in the SC following visual training may be a result of two phenomena: (1) as a result of changes to retinotectal synapses independently of the V1; (2) or enhancement in the V1 that modulates response in the SC. To resolve this problem we selectively inactivated the V1 through the application of xylocaine solution on the surface of the V1 for the duration of visual training. Chemical inactivation of the V1 caused strong silencing of cortical responses during visual training (Fig 4A, B). The chemical inactivation prevented cortex from response enhancement as presented in Figure 4C and revealed no difference in amplitude before and after training (83 ± 4 %, n = 14, p = 0.25). This result confirms an effective deactivation of the V1 during training and blocking the learning effect in this structure. The comparison between pre- and post-training recordings in the SC showed significant enhancement of the response as a result of the visual training while V1 deactivation. In the SGS layer, the amplitude of the visual response was significantly greater in post-training control compared to pre-training recording (Fig. 4C; 212 ± 26 %, n = 10, p = 0.0025). Moreover, the amplitude was higher than changes evoked in the SO layers (Fig. 4C; 161 ± 23 %, n = 10, p = 0.03). We observed an increase of the collicular VEP amplitudes during visual training even when cortical activity was silenced (for SGS; F(3,9) = 6.59, p = 0.005, for SO; F(3,9) = 9.28, p = 0.02). For the SGS layers (Fig. 4D) VEP amplitude was the highest after 3 hours of visual training compared to control time (262 ± 36 %, p = 0.01), then 2^nd^ hour (222 ± 25 %, p = 0.02) and after 1^st^ hour (152 ± 10 %, p = 0.08). We found similar tendency in the SO layer (Fig. 4E) where the potentiation of visual response was the greatest after 3^rd^ hour of stimulation (222 ± 31 %, p = 0.04), then 2^nd^ hour (213 ± 36 %, p = 0.1) and after 1^st^ hour (173 ± 24 %, p = 0.1).

**Figure 4.**
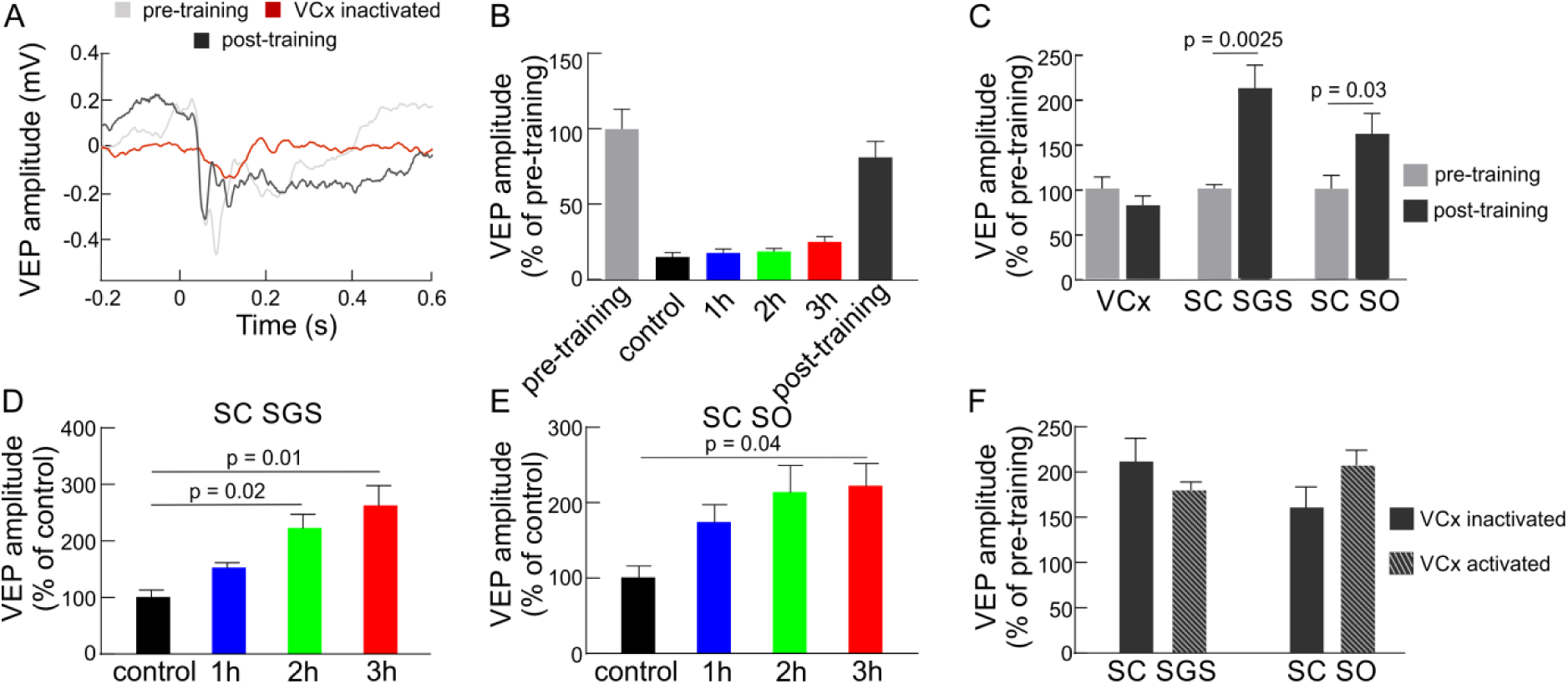
Cortical inactivation does not affect plasticity in the SC. A, V1 VEP from the single experiment before (gray line), during visual training when xylocaine was applied (red line), and after training when xylocaine was washed out (black line). B, Silencing of cortical responses during visual training as a result of xylocaine application. C, Comparison of average responses before (gray) and after (black) visual training in the V1, SGS and SO layers of the SC when cortical activity was suppressed. D, E, Average response amplitudes in the SC (SGS and SO, respectively) for every hour of the visual training while V1 silencing. F, Comparison of mean values of post-training VEP amplitudes in the SC between two experimental conditions: with V1 active (n = 10) and V1 inactive (n = 10).

A comparison of the visual training effects of the two conditions (V1 activated and inactivated) did not show a significant difference in the SC (Fig. 4F). The increase of visual response for every hour of training in both layers was similar in both V1 conditions. This result indicates that the temporary blocking of V1 did not inhibit VEP plasticity in SC evoked by repeated visual training. We can also conclude that the enhancement of VEP amplitude in the SC occurred mainly through a retinotectal synapse.

## Discussion

Our results showed that the 3 hours visual training evoked strong enhancement of the visually evoked potentials both in the V1 and the SC. Moreover, our paradigm of repetitive visual training evoked a stronger increase in the response in the SC than in the V1 (Fig. 2). We confirm that visual training causes enhancement of VEP amplitude in the V1 as it was described before [7,9–12,17] and extends the knowledge showing positive results of this training in the rat’s SC. Specifically, we showed in an electrophysiological study that single training session, lasting several hours, induces plasticity of visual responses.

Based on the literature we can enumerate several well-described paradigms of repetitive visual stimulation which differ from each other mainly in the type of visual stimulus, presentation timing, number of repetitions and the frequency of the stimulus [9,11,13–15,28]. One well–known protocol uses repeated presentations of a specifically oriented visual stimulus through several days that causes stimulus-selective potentiation of a cortical response in awake mice and enhancement of a signal detection power in humans [29]. This type of plasticity is well described in layer IV of the V1 [9,12]. In our study, we found that the potentiation of the visual response to a flash stimulus also occurs in the infragranular layers. Specific types of repeated visual stimulation (shorter and more intense) are also able to induce the modulation of VEP plasticity and modifications of synaptic connectivity in the mature V1. Studies carried out on humans demonstrated how 10 min presentation of checkerboard reversals [8] resulted in sustained amplitude modulation of early components of subsequent VEPs, whereas a rapid (9 Hz, for 2 min) checkerboard stimulation might induce the enhancement of visual response in the adult rats V1 [15]. Our experimental paradigm provides a form of rapid visual training consisted of a series of flashes repeated every 15 minutes through 3 hours. The frequency of stimulus presentation (0.5 Hz) is lower than was used by Clapp et al. [15] nevertheless it was sufficient to evoke significantly enhanced responses in the visual cortex and superior colliculus.

Flashing stimuli compared to drifting sinusoidal gratings or reversal checkerboards are rather strong stimuli, thus can evoke changes faster. It was shown before that in adult mice flashing stimuli robustly evoke long-term changes in the V1 neuronal response and increase the broadband power of LFP signal [30,31]. Thus, the flash stimulus might be successfully used for the induction of modulation in the neuronal response [32], which we confirmed in our study.

It is considered that the reinforcement of neuronal response occurring following repetitive stimulation in the visual system in awake animals, including humans might be triggered by increasing the number or gain of neurons involved in the response to the trained stimulus [14,28]. In our study, we observed facilitation of visual response to flash stimulus although the experiments were carried out on anesthetized rats. Previous studies also confirmed the occurrence of learning processes in the visual system in animals under deep anesthesia, where LTP dependent of NMDA receptors was effectively induced in the V1 through electrical theta-burst stimulation of the visual pathway [6,13] or via repetitive visual stimulation [15].

We found that 3 hours of repeated visual stimulation evoked enhancement not only in the cortex but also in the SC (Figs. 2 and 3). We also show for the first time that, stronger response enhancement occurred in the SC than in the V1. So far, little attention has been devoted to the investigation of this effect. Zhang et al. [23] shown that repetitive exposure to dimming stimuli effectively induced the LTP of developing retinotectal synapses in *Xenopus* tadpoles. This effect is attributed mainly to changes in synaptic efficacy at retinotectal projections. However, potentiation of neuronal response at this level of the visual processing can derive also from the cortex due to inputs from layer 5 of the V1 [33,34]. Our results proved that collicular response potentiation is not changed when V1 is blocked during repetitive visual training. This indicates that the increase of neuronal responses in the SC is most likely due to the enhancement of the retinotectal projection. There are supportive studies performed on rats where ablation of the V1 caused facilitation of the LTP formation in the SC indicating a suppressive influence of the V1 inputs into the SC [35,36]. In our studies, we observed potentiation of the SC response during visual training regardless of whether the V1 was activated or inactivated (Fig. 3 and 4).

In summary, the data presented here show a new form of plasticity occurring after 3 hours of repeated visual training in the primary visual cortex and superior colliculus. We proved that the enhancement of neuronal responses in the SC following our paradigm of visual stimulation occurred independently of the V1, most likely through retinotectal projection. Further research will be needed to better understand the mechanisms responsible for this phenomenon.

## Acknowledgments

This work was funded by the National Science Centre Poland Grant 2017/25/N/NZ4/02914. We dedicate this work to Prof. Wioletta Waleszczyk, our friend, mentor and supervisor.

